# ANNOgesic: A Swiss army knife for the RNA-Seq based annotation of bacterial/archaeal genomes

**DOI:** 10.1101/143081

**Authors:** Sung-Huan Yu, Jörg Vogel, Konrad U. Förstner

## Abstract

To understand the gene regulation of an organism of interest, a comprehensive genome annotation is essential. While some features, such as coding sequences, can be computationally predicted with high accuracy based purely on the genomic sequence, others, such as promoter elements or non-coding RNAs are harder to detect. RNA-Seq has proven to be an efficient method to identify these genomic features and to improve genome annotations. However, processing and integrating RNA-Seq data in order to generate high-resolution annotations is challenging, time consuming and requires numerous different steps. We have constructed a powerful and modular tool called ANNOgesic that provides the required analyses and simplifies RNA-Seq-based bacterial and archaeal genome annotation. It can integrate data from conventional RNA-Seq and dRNA-Seq, predicts and annotates numerous features, including small non-coding RNAs, with high precision. The software is available under an open source license (ISCL) at https://pypi.org/project/ANNOgesic/.

## 1 Background

As the number of available genome sequences has rapidly expanded in the data bases, numerous tools have been developed that can detect genomic features of interest based on the genome sequence. Prominent representatives are Glimmer to identify open reading frames (ORFs) [13], tRNAscan-SE [56] to spot tRNAs and RNAmmer to find rRNAs [35]. Pipelines like Prokka [57] or ConsPred [68] combine such tools and are able to search multiple features in bacterial and archaeal genomes. Still, these tools that make their predictions purely based on the genome sequences can predict features like transcriptional start sites and non-coding RNAs, if at all, only with low confidence.

Recent developments in high-throughput sequencing offer solutions to this problem. RNA-Seq has revolutionized how differential gene expression can be measured and is widely used for this purpose [47]. Besides this it has also been applied in numerous cases to improve the genome annotation of bacteria [3,6,58], archaea [69] and eukaryotes [23]. RNA sequencing based protocols like differential RNA-Seq (dRNA-Seq) [4, 59], Term-seq [12] and ribosome profiling [29, 66] have been applied to globally detect transcriptional start sites (TSSs), small non-coding RNAs (ncRNAs/sRNAs), terminators, ORFs and sRNA regulatory target but require dedicated data processing. While there are tools that can process RNA-Seq data in order to predict genome-wide features like TSSs based on dRNA-Seq data [1,16,30] or based on conventional RNA-Seq data [42, 54], there has been, so far, no solution that combines different predictions of genomic features and compiles them into a consistent annotation.

Here we present ANNOgesic - a modular, command-line tool that can integrate data from different RNA-Seq protocols like dRNA-Seq as well as conventional RNA-Seq performed after transcript fragmentation and generate high-quality genome annotations that include features missing in most bacterial annotations (e.g. small non-coding RNAs, UTRs, TSSs, operons). The central approach is to detect transcript boundaries and then subsequently attach further information about type as well as function to the predicted features and also to infer interactions between them. Several of ANNOgesic’s core functions represent new implementations that are not found in other programs. Those are accompanied by third-party tools that are embedded into ANNOgesic and are due to that accessible via a consistent command-line interface. Furthermore their results are improved e.g. by dynamic parameter-optimizations or by removing false positives. Numerous visualizations and statistics help the user to quickly evaluate the feature predictions. The tool is modular and was intensively tested with several RNA-Seq data sets from bacterial as well as from archaeal species.

## 2 Materials and Methods

### 2.1 Modules of ANNOgesic

ANNOgesic consists of the following modules - their names indicate their functions: Sequence modification, Annotation transfer, SNP/Mutation, Transcript, TSS, Terminator, UTR, Processing site (PS), Promoter, Operon, sRNA, sRNA target, sORF, GO term, Protein-protein interaction network, Subcellular localization, Riboswitch, RNA thermometer, Circular RNA, and CRISPR. Several potential workflows connecting these modules are displayed in Figure S1. An overview of the novelties and improvements of the modules in ANNOgesic are listed in Table S1.

Depending on the task to one wish to perform, ANNOgesic requires a specific set of input information - either as coverage information in wiggle format or alignments in BAM format. This can be generated by short read aligners like STAR [15], segemehl [27], or a full RNA-Seq pipeline like READemption [18]. Certain modules additionally require annotations in GFF3 format. In case a sufficient genome annotation is not available, ANNOgesic can perform an annotation transfer from a closely related strain.

### 2.2 Implementation and installation

ANNOgesic’s source code is implemented in Python 3 and hosted at https://pypi.org/project/ANNOgesic/. The comprehensive documentation can be found at http://annogesic.readthedocs.io/ and releases are automatically submitted to Zenodo (https://zenodo.org/) to guarantee a long term availability. It can be easily installed using pip (https://pip.pypa.io). In order to provide a frictionless installation including non-Python dependencies, we additionally offer a Docker image (https://hub.docker.com/r/silasysh/annogesic/) [44].

### 2.3 Optimization of the parameter set for TSSpredator

For several parts of ANNOgesic the selection of parameters has a strong impact on the final results. Especially the TSS prediction – building on TSSpredator [16] – requires a sophisticated fine-tuning of several parameters (namely height, height reduction, factor, factor reduction, enrichment factor, processing site factor and base height). To overcome the hard task of manual parameter selection, ANNOgesic optimized the parameters by applying a genetic algorithm, a machine learning approach, [21] which is trained based on a small user curated set of TSS predictions. This approach has the advantage of being able to find global, not only local, optima. The process of optimization is composed of three parts - random change, large change, and small change (Figure 1). In this context, a global change means a random allocation of values to all parameters, a large change is a random allocation of values to two parameters, while a small change is adding or subtracting a small fraction to or from one parameter value. The result of each iteration is evaluated by a decision statement (Equation 1).

**Figure 1.**
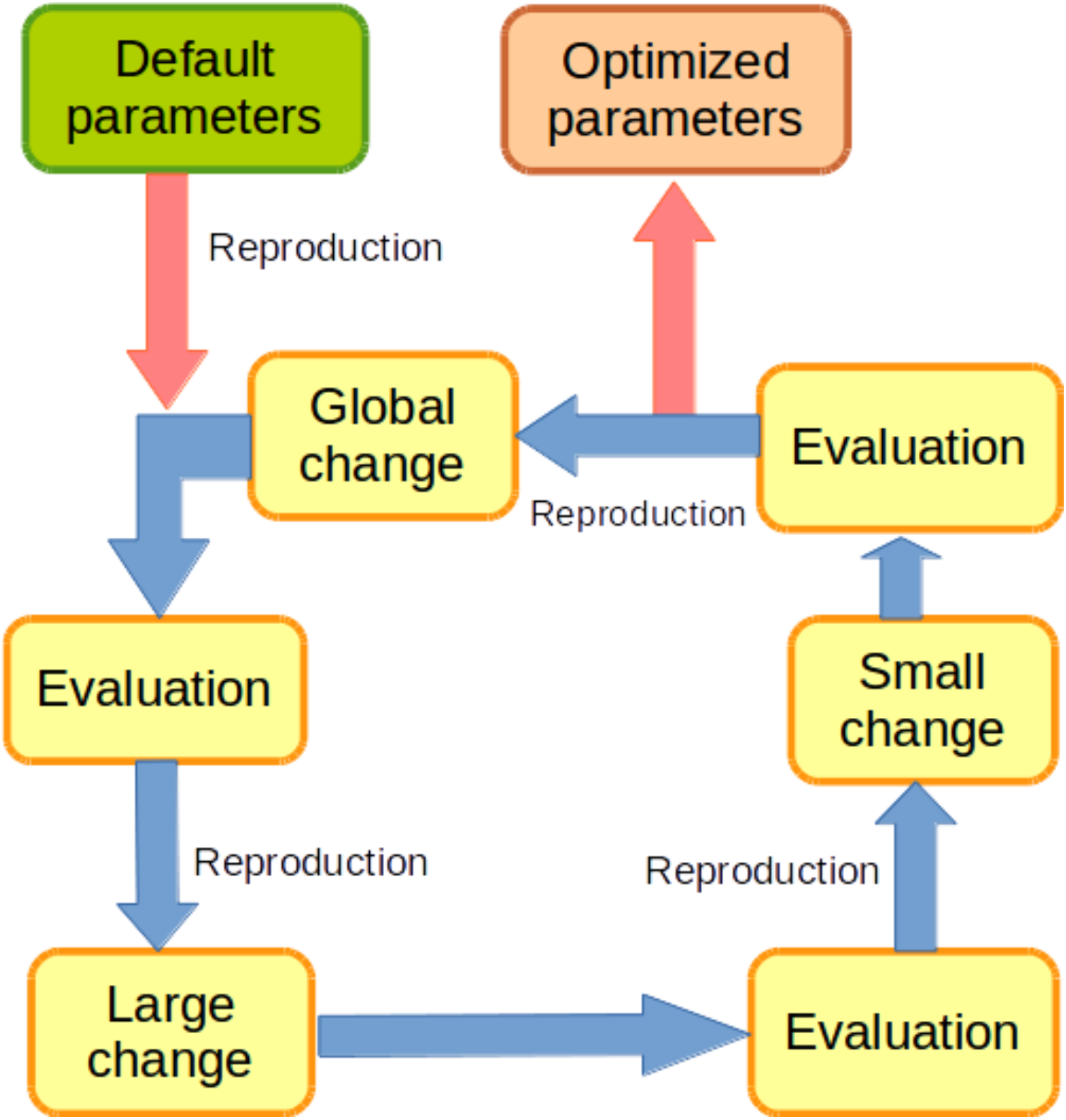
Schema of the genetic algorithm for optimizing the parameters of TSSpredator. It starts from the default parameters. These parameter sets will go through three steps - global change (change every parameter randomly), large change (change two of the parameters randomly), and then small change (adds/subtracts a small fraction to one of the parameters). It will then select the best parameter set for reproduction when one step is done. Usually, ANNOgesic can achieve the optimized parameters within 4000 runs.

### 2.4 Test data sets

In order to test ANNOgesic’s performance, we applied it to RNA-Seq data sets originating from *Helicobacter pylori* 26695 [4, 58] and *Campylobacter jejuni* 81116 [16]. The dRNA-Seq data sets were retrieved from NCBI GEO where they are stored under the accession numbers GSE67564 and GSE38883, respectively. For *H. pylori* conventional RNA-Seq data – i.e. without TEX treatment (which degrades transcripts without a 5’-triphosphate) and with fragmentation of the transcript before the library preparation – was also retrieved from NCBI SRA (accession number SRR031126). Moreover, for assessing the performance of ANNOgesic, dRNA-Seq and conventional RNA-Seq data sets of *Escherichia coli* K12 MG1655 were downloaded from NCBI GEO (accession number: GSE55199 and GSE45443 (only the data of the wild type strain was retrieved) [42,65]. The ANNOgesic predictions generated using these data sets of *Escherichia coli* K12 MG1655 were compared to the databases RegulonDB, EcoCyc and DOOR^2^ [20,22,24,31,41,51].

## 3 Results

### 3.1 Correction of genome sequences and annotations

All genomic features that can be detected by ANNOgesic are listed in Table 1. In order to demonstrate and test ANNOgesic’s performance we have analyzed RNA-Seq data of *H. pylori* 26695 and *C. jejuni* 81116 and discuss the prediction results as examples in the following sections.

**Table 1.** Overview of feature predictions for *H. pylori* 26695 and *C. jejuni* 81116

|  |  | *H. pylori* 26695 | *C. jejuni* 81116 |
| --- | --- | --- | --- |
| Gene |  | 1560 | 1685 |
| CDS | Total | 1448 | 1630 |
|  | Expressed | 1406 | 1513 |
| Transcript |  | 1716 | 1147 |
| TSS | Total | 2458 | 1242 |
|  | Primary | 703 | 565 |
|  | Secondary | 156 | 92 |
|  | Internal | 719 | 360 |
|  | Antisense | 1161 | 510 |
|  | Orphan | 111 | 30 |
| Processing site |  | 281 | 345 |
| Terminator | Total | 820, (437) | 874, (375) |
|  | TransTermHP | 631, (314) | 655, (269) |
|  | Convergent genes | 229, (151) | 276, (145) |
| UTR | 5' UTR | 693 | 560 |
|  | 3' UTR | 325 | 286 |
| sRNA | Total | 184 | 40 |
|  | Intergenic | 60 | 16 |
|  | Antisense | 85 | 21 |
|  | 5' UTR-derived | 10 | 0 |
|  | 3' UTR-derived | 23 | 2 |
|  | InterCDS-derived | 6 | 1 |
| Operon | Total | 1716 | 1147 |
|  | Monocistronic | 269 | 386 |
|  | Polycistronic | 285 | 324 |
|  | No CDS associated | 1162 | 437 |
| sORF |  | 150 | 25 |
| Riboswitch |  | 11 | 14 |
| RNA thermometer |  | 4 | 8 |
| circular RNA |  | 0 | 1 |
| CRISPR |  | 0 | 1, (8) |

The numbers in brackets for terminator and CRISPR represent occurrences of terminators with coverage drop and repeat units of CRISPR, respectively. For the prediction of terminators ANNOgesic only keeps the high confidence ones in case a CDS is associated with multiple terminators.

#### 3.1.1 Genome sequence improvement and SNP/mutation calling

Conventionally, differences in the genome sequence of a strain of interest and the reference strain are determined by DNA sequencing. However, RNA-Seq reads can also be re-purposed to detect such SNPs or mutations that occur in transcribed regions which can help to save the resources required for dedicated DNA sequencing or DNA SNP microarray measurements. The two drawbacks of this method are that only locations which are expressed can be analyzed and that, due to RNA editing, changes could be present only in the RNA level and are not found in the genome. On the other hand, it has been shown to be a valid approach for eukaryotic species and that the majority of SNPs are found in the expressed transcripts [10,11]. In conclusion, such an analysis could be useful to generate hypotheses that then need to be tested with complementary methods. ANNOgesic offers the user to perform the SNP/mutation calling via SAMtools [36] and BCFtools [36] applying read counting-based filtering.

### 3.1.2 Annotation transfer

ANNOgesic integrates RATT [50], which can detect the shared synteny and mutations between a reference and query genome to transfer annotation (i.e. genes, CDSs, tRNAs, rRNAs) by applying MUMmer [34]. For the chosen strains, *H. pylori* 26695 and *C. jejuni* 81116 annotation files in GFF3 format were be obtained from NCBI RefSeq. Due to this there was no need to transfer the annotation from a closely related strain.

### 3.2 Detection of transcripts

Knowing the exact boundaries and sequence of a transcript is crucial for a comprehensive understanding of its behaviour and function. For example, UTRs can be the target of regulation by sRNAs or small molecules (e.g. riboswitches) [7,67] or even sources of sRNAs [9]. Unfortunately, most bacterial annotations only cover the protein coding regions while the information about TSSs, terminators and UTRs is lacking. To address this issue, ANNOgesic combines several feature predictions for a reliable detection of transcripts and their boundaries (Figure 2).

**Figure 2.**
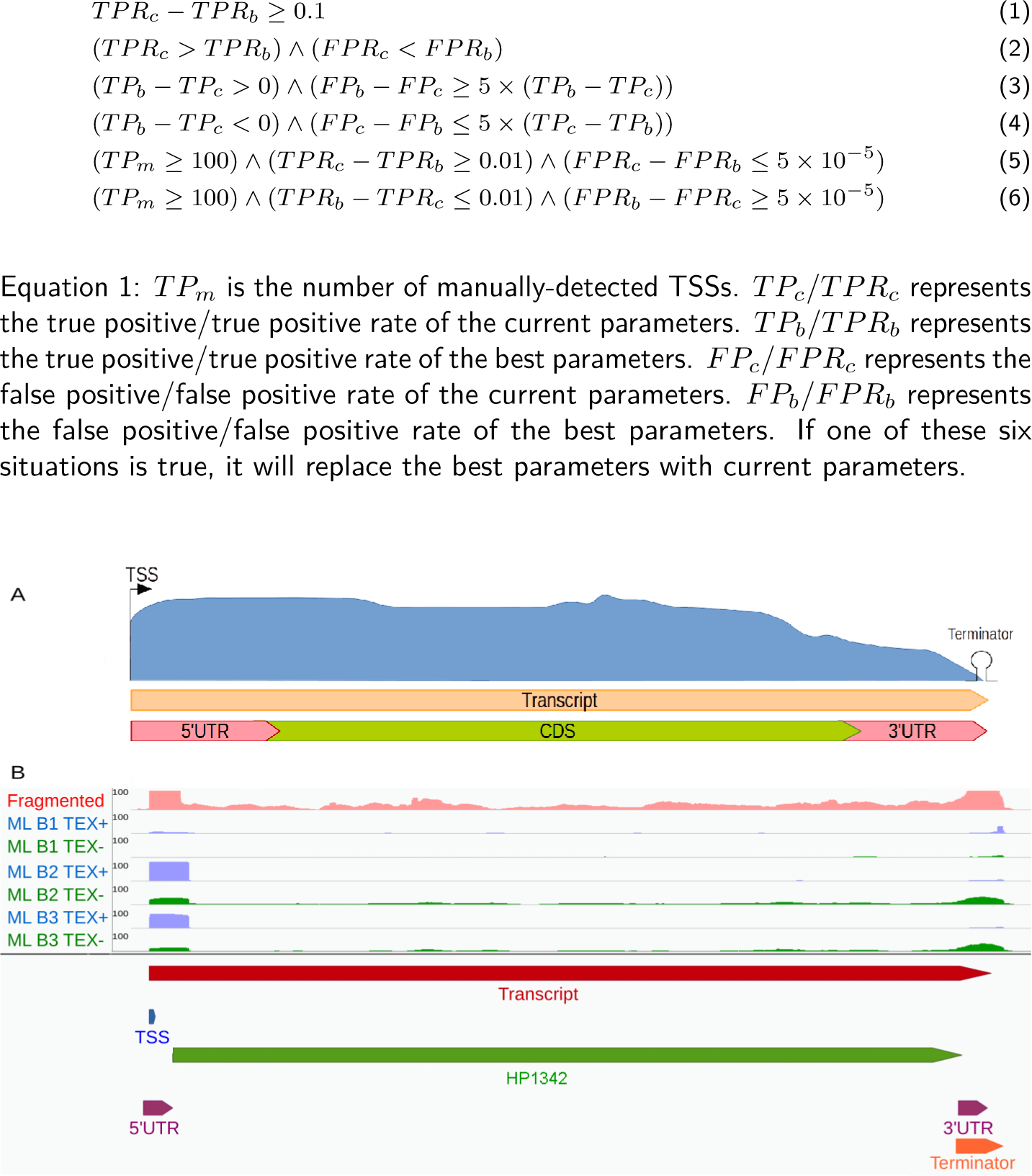
Transcript boundary detection. **(A)** Schema: ANNOgesic can predict TSSs, terminators, transcripts, genes and UTRs and integrate them into a comprehensive annotations. **(B)** Gene HP1342 of *H. pylori* 26695 as an example. The pink coverage plot represents RNA-Seq data of libraries after fragmentation, the blue coverage plots TEX+ libraries of dRNA-Seq, the green coverage plots TEX- libraries of dRNA-Seq. Transcript, TSS, terminator, and CDS are presented as red, blue, orange, and green bars, respectively. The figure shows how the transcript covers the whole gene location and how UTRs (presented by purple bars) can be detected based on the TSS, transcript, terminator, and gene annotations.

#### 3.2.1 Coverage based transcript detection

There are numerous tools available for the detection of transcripts (e.g. [17]), but most of them are optimized for the assembly of eukaryotic transcripts. Due to this, we combined several heuristics to perform such predictions. Nucleotide coverage data is used for define the expressed regions, and genome annotations are applied for extending or merging the gene expressed regions to form complete transcripts. Several parameters like the threshold of coverage values can be set by the user to fine-tune the predictions (Figure 3).

**Figure 3.**
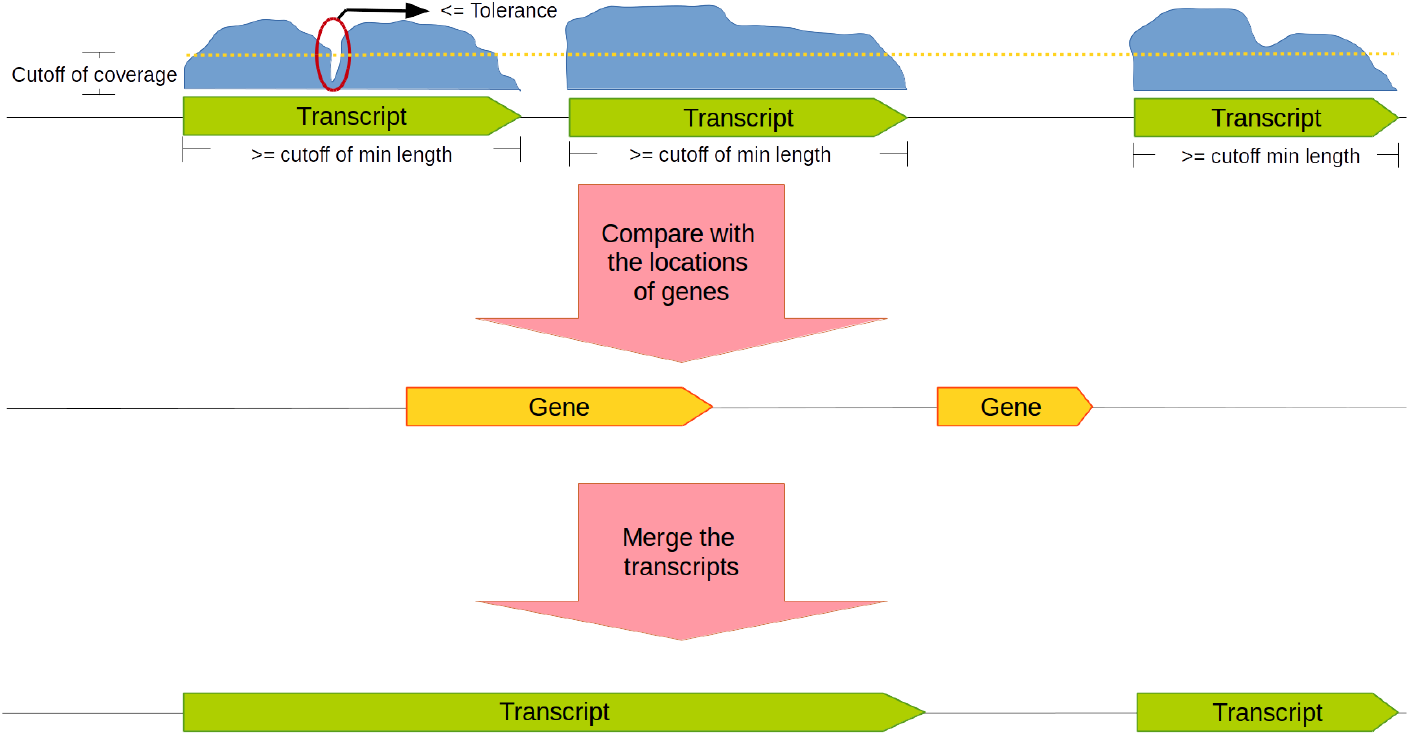
Coverage-based transcript detection. If the coverage (blue curve-blocks) is higher than a given coverage cut-off value (dashed line) a transcript will be called. The user can set a tolerance value (i.e. a number of nucleotides with a coverage below the cut-off) on which basis gapped transcripts are merged or are kept separated. Information of gene positions can also be used to merge transcripts in case two of them overlap with the same gene.

By running ANNOgesic’s subcommand for transcript prediction, we detected 1715 transcripts in *H. pylori* 26695 and 1147 transcripts in *C. jejuni* 81116. These transcripts cover 1520 and 1568 genes which shows that 97% and 93% of the known genes are expressed in at least one condition, respectively.

#### 3.2.2 Transcriptional start sites

For the prediction of TSSs, ANNOgesic builds on TSSpredator [16], which takes dRNA-Seq coverage data as input. The outcome of TSSpredator’s predictions depends strongly on the setting of numerous parameters and fine-tuning those can be time consuming. Due to this, a parameter optimization was implemented in ANNOgesic that builds on a small, manually curated set of TSSs to find optimal values.

In order to test the performance of ANNOgesic, we manually annotated TSSs in the first 200 kb of the genome of *H. pylori* 26695 and *C. jejuni* 81116 (Table S4 and S5). This set was used to benchmark the prediction of TSSpredator with default settings as well with the parameters optimized by ANNOgesic. For the test set, we manually annotated TSSs from first 200 kb to 400 kb in the genome of *H. pylori* 26695 and *C. jejuni* 81116 (Table S4 and S5). As displayed in Table 2, the optimization had minor sensitivity improvements in *H. pylori* 26695 (from 96.8% to 99.6%), while strongly increased the sensitivity for the TSS prediction for *C. jejuni* 81116 (67.1% to 98.7%) while keeping the same level of specificity. To underpin those findings, we looked at the overlap of the predicted TSS and predicted transcripts. This was nearly the same for *H. pylori* 26695 (82% for default and 83% for optimized setting) but also increased significantly for *C. jejuni* 81116 from 81% for default parameters to 96% with optimized parameters.

Moreover, TSSs are classified depending on their relative positions to genes by TSSpredator. Based on these classifications, Venn diagrams representing the different TSS classes are automatically generated (Figure S2).

#### 3.2.3 Processing sites

Several transcripts undergo processing, which influences their biological activity [9,25]. In order to detect PSs based on dRNA-Seq data, ANNOgesic facilitates the same approach as described for TSS detection but searches for the reverse enrichment pattern (i.e. a relative enrichment in the library not treated with TEX). As done for the TSSs, we manually annotated the PSs in the first 200 kb of the genomes. Based on these manually curated sets, we performed parameter optimization on the test set (manually-curated from first 200 kb to 400 kb, Table S6 and S7, Table 2) and could improve the prediction of PSs by TSSpredator [16]. With optimized parameters 281 and 345 PSs were detected in *H. pylori* 26695 and *C. jejuni* 81116, respectively.

**Table 2.** Comparison of default and optimized parameters of TSSpredator for TSS and PS prediction

| Strains | Parameters | Sensitivity (TP) | Specificity (FP) |
| --- | --- | --- | --- |
| TSS |  |  |  |
| *H. pylori* 26695 | Default | 96.8% (244) | 99.98% (32) |
|  | Optimization | 99.6% (251) | 99.98% (32) |
| *C. jejuni* 81116 | Default | 67.1% (104) | 99.98% (31) |
|  | Optimization | 98.7% (153) | 99.99% (7) |
| PS |  |  |  |
| *H. pylori* 26695 | Default | 92.9% (26) | 99.99% (7) |
|  | Optimization | 92.9% (26) | 99.99% (7) |
| *C. jejuni* 81116 | Default | 61.3% (19) | 99.99% (2) |
|  | Optimization | 93.5% (29) | 99.99% (6) |

The numbers in brackets represent true positive or false positive.

#### 3.2.4 *ρ*-independent terminators

While the TSSs are in general clearly defined borders, the 3‘-end of a transcript is often not very sharp. A commonly used tool for the prediction of the 3’-end of a transcript is TransTermHP [33], which detects *ρ*-independent terminators based on genome sequences. Manual inspection showed us that TransTermHP predictions are not always supported by the RNA-data (Figure S3E and S3F). This could be due to the lack of expression in the chosen conditions. Additionally, certain locations in 3’-ends that may be *ρ*-independent were not detected by TransTermHP. Due to this, we extended the prediction by two further approaches based on RNA-Seq coverage and the given genome sequence. At first, terminators predicted by TransTermHP that show a significant decrease of coverage are marked as high-confidence terminators. For this, the drop of coverage inside the predicted terminator region plus 30 nucleotides up and downstream is considered as sufficient if the ratio of the lowest coverage value and the highest coverage value is at a user-defined value (the default value is 0.5, and the schemes and examples are shown in Figure S3). In order to improve the sensitivity, an additional heuristic for the detection of *ρ*-independent terminators was developed. In this approach, only converging gene pairs (i.e. the 3’-end are facing to each other) are taken into account (Figure S4). In case the region between the two genes is A/T-rich and a stem-loop can be predicted in there, the existence of a *ρ*-independent terminator is assumed. As default, the region should consist of 80 or less nucleotides, the T-rich region should contain more than 5 Thymines, the stem-loop needs to be 4 - 20 nucleotides, the length of the loop needs to be between 3 and 10 nucleotides and maximum 25% of the nucleotides in the stem should be unpaired.

#### 3.2.5 UTRs

Based on the CDS locations and the above described detection of TSSs, terminators and transcripts, 5‘ UTR and 3’ UTR can be annotated by ANNOgesic. Additionally, it visualizes the distribution of UTR lengths in a histogram (as shown in Figure S5).

#### 3.2.6 Promoters

ANNOgesic integrates MEME [2] which detects ungapped motifs and GLAM2 [19] which discovers gapped motifs for the detection and visualization of promoter motifs. The user can define the number of nucleotides upstream of TSSs that should be screened and the length of potential promoter motifs. The motifs can be generated globally or for the different types of TSSs (example in Figure S6).

#### 3.2.7 Operons

Based on the TSS and transcript prediction, ANNOgesic can generate statements regarding the organization of genes in operons and suboperons as well as report the number of monocistronic operons and polycistronic operons (Figure 4).

**Figure 4.**
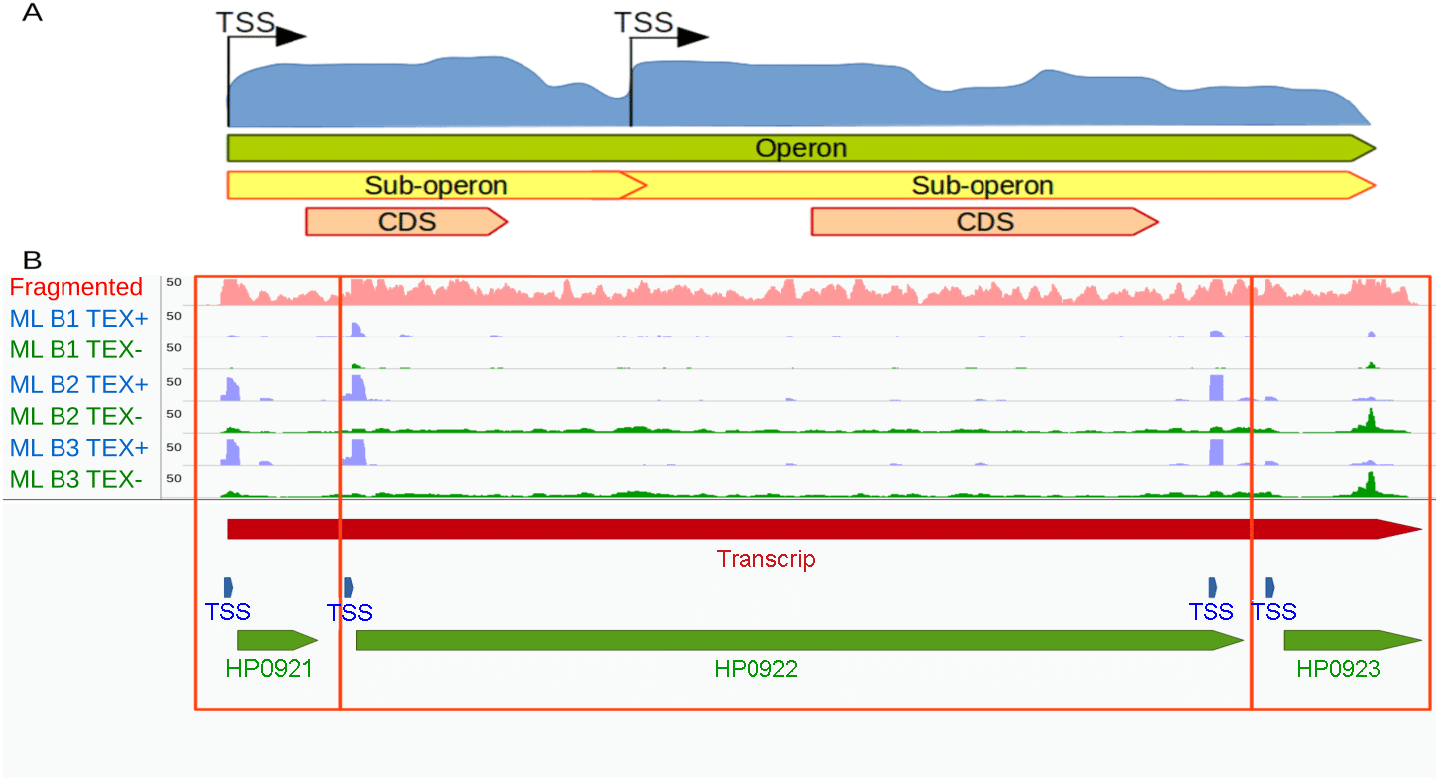
Operon and sub-operon detection. **(A)** If there are more than one TSSs which does not overlap with genes located within one operon, the operon can be divided to several sub-operons based on these TSSs. **(B)** An example from *H. pylori* 26695. The coverage of RNA-Seq with fragmentation, TEX+ and TEX-of dRNA-Seq are shown in pink, blue and green coverages, respectively. TSSs, transcripts/operons and genes are presented as blue, red and green bars, respectively. The two genes are located in the same operon, but also in different sub-operons (two empty red squares).

### 3.3 Detection of sRNAs and their targets

The detection of sRNAs based on RNA-Seq data is a non-trivial task. While numerous sRNAs are found in intergenic regions, several cases of 5‘ and 3’ UTR-derived sRNAs are reported [9,28,45,60]. ANNOgesic offers the detection of all classes, combined with a detailed characterization of the sRNA candidates (Figure 5).

**Figure 5.**
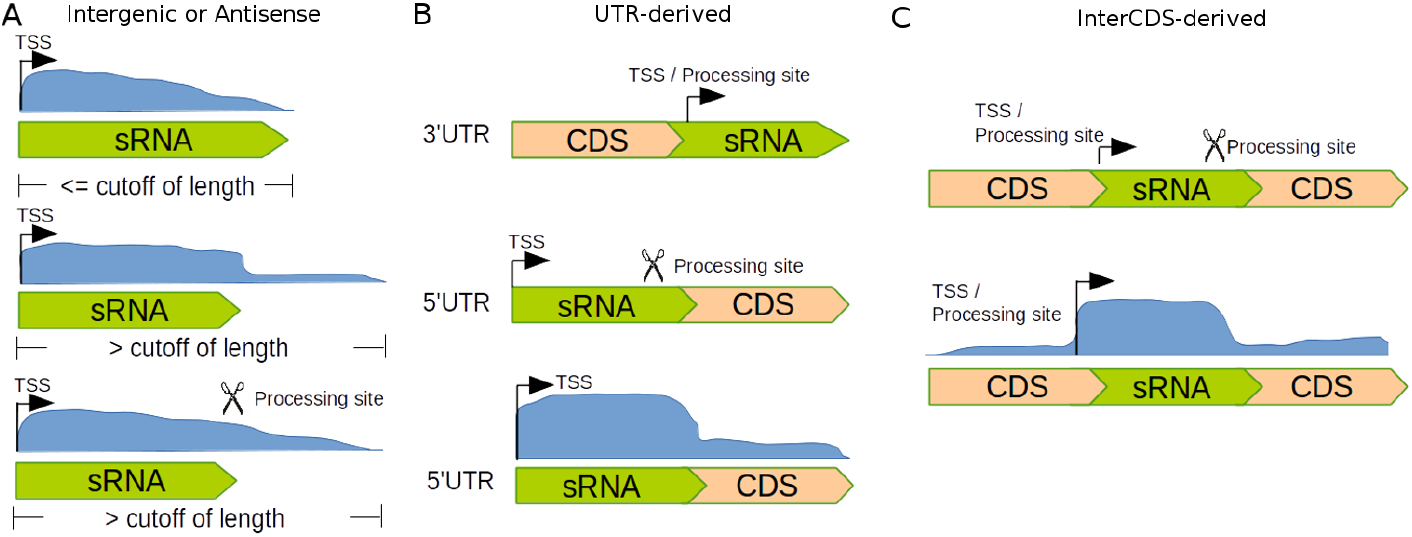
Detection of intergenic, antisense and UTR-derived sRNAs. Detection of intergenic, antisense and UTR-derived sRNAs. The length of potential sRNAs should be within a given range and their coverages should exceed a given cut-off coverage. **(A)** Detection of intergenic and antisense sRNAs. There are three cases shown: In the upper panel the transcript starts with a TSS, and length of the transcript is within the expected length. In the one in the middle the transcript starts with a TSS but the transcript is longer than an average sRNA; in that case ANNOgesic will search the coverage (blue region) for a point at which the coverage is decreasing rapidly. The image at the bottom is similar to the one in the middle, but the sRNA ends with a PS. **(B)** Detection of UTR-derived sRNAs. 3‘ UTR-derived sRNAs: if the transcript starts with a TSS or PS, it will be tagged as a 3’ UTR-derived sRNA. For 5’ UTR-derived sRNAs: if the transcript starts with a TSS and ends with a PS or the point where the coverage significant drops. **(C)** Detection of interCDS-derived sRNAs: Similar to the 5’ UTR-derived approach but the transcript starts with a PS.

In order to classify newly detected intergenic transcripts as sRNAs, ANNOgesic tests several of their features. If a BLAST+ [8] search of a transcript finds homologous sequences in BSRD [37] – a database that stores experimentally confirmed sRNAs – the transcript gets the status of an sRNA. The user can also choose further databases for searching homologous sequences. In case a search against the NCBI non-redundant protein database leads to a hit it is marked as potentially protein-coding. Otherwise, a transcript must have a predicted TSS, form a stable secondary structure (i.e. the folding energy change calculated with RNAfold from Vienna RNA package [38] must be below a user defined value) and their length should be in the range of 30 to 500 nt in order to be tagged as an sRNA. All these requirements are used per default but can be modified or removed via ANNOgesic’s command line parameters. ANNOgesic stores the results of all analyses and generates GFF3 files, fasta files, secondary structural figures, dot plots, as well as mountain plots based on those predictions.

For sRNAs that share a transcript with CDSs – 5‘ UTR, inter-CDS, or 3’ UTR located sRNAs – we implemented several detection heuristics (Figure 5B / C). 5‘ UTR-derived sRNAs must start with a TSS and show a sharp drop of coverage or a PS in the 3’-end. The requirement for the detection of inter-CDS located sRNAs is either a TSS or a PS as well as a coverage drop at the 3’-end or a PS. Small RNAs derived from the 3’ UTR are expected to have a TSS or a PS and either end with the transcript or at a PS. After the detection of a *bona fide* sRNA, the above described quality filters (length range, secondary structure etc.) are applied to judge the potential of a candidate (examples are shown in Figure S7, S8). For the validation of sRNA candidates in our test case, the described sRNAs of two publications were chosen. Sharma *et al*. [58] described 63 sRNAs of which 4 were not expressed in the condition of the test data set (removed from the dataset) (Figure S9). Of these 59 53 (90%) were detected by ANNOgesic. In the *C. jejuni* 81116 set, 31 sRNAs were described by Dugar *et al*. [16], and ANNOgesic could recover 26 (84%). The sRNA ranking system provided by ANNOgesic is displayed in Figure S10 and Equation S1.

In order to deduce potential regulatory functions of newly-predicted sRNAs, ANNOgesic performs prediction of interaction between them and mRNAs using RNAplex [38, 63], RNAup [38,46], and IntaRNA [40]. The user can choose if only interactions supported by all tools are reported.

### 3.4 Detection of sORFs

All newly detected transcripts that do not contain a previously described CDS as well all 5‘ UTRs and 3’ UTRs are scanned for potential sORFs [61] (Figure 6). For this, ANNOgesic searches for start and stop codons (non-canonical start codons are not included, but can be assigned by the user) that constitute potential ORFs of 30 to 150 base-pairs. Furthermore, ribosomal binding sites (based on the Shine-Dalgarno sequence, but different sequences can be assigned as well) between the TSS and 3 to 15 bp upstream of the start codon are required for a *bona fide* sORF.

**Figure 6.**
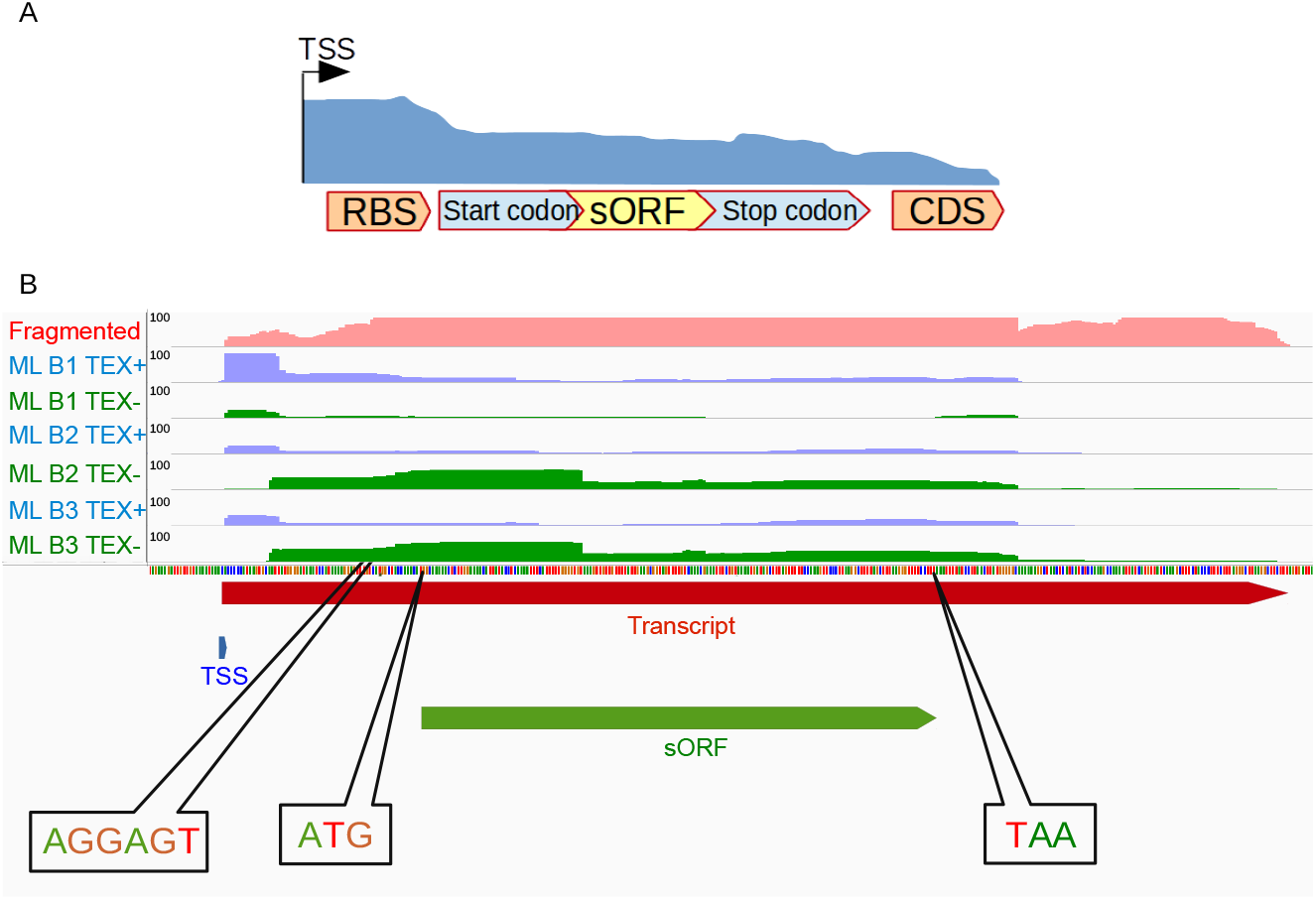
sORF detection. **(A)** An sORF must contain start codon and stop codon within transcript and should be inside of a given length range (default 30 - 150 nt). Additionally, a ribosomal binding site must be detected between the TSS and the start codon. **(B)** An example from *H. pylori* 26695. The coverage of RNA-Seq (fragmented libraries), TEX+ and TEX- (dRNA-Seq) are shown as pink, blue and green coverages, respectively. The TSS, transcript and sORF are presented as blue, red and green bars, respectively.

### 3.5 Detection of functional related attributes

In order to facilitate a better understanding of the biological function of known and newly detected transcripts, ANNOgesic predicts several attributes for these features.

This includes the allocation of GO as well as GOslim [64] terms to CDSs via searching of protein ids in Uniprot [39]. The occurrence of groups is visualized for expressed and non-expressed CDSs (Figure S11). Furthermore the subcellular localization is predicted by PSORTb [70] for the proteins (Figure S12). Additionally, the protein entries are enriched by protein-protein interaction information retrieved from STRING [62] and PIE [32] (examples in Figure S13).

### 3.6 Circular RNAs

ANNOgesic integrates the tool “testrealign.x” from the segemehl package for the detection of circular RNAs [26] and adds a filter to reduce the number of false positive. Candidates for circular RNAs must be located in intergenic regions and exceed a given number of reads.

### 3.7 CRISPRs

CRISPR/Cas systems represent a bacterial defence system against phages and consist of repeat units and spacers sequences as well as Cas proteins [55]. The CRISPR Recognition Tool (CRT) [5] is integrated into ANNOgesic and extended by comparison of CRISPR/Cas candidates to other annotations to remove false positive (Figure S14).

### 3.8 Riboswitches and RNA thermometers

Riboswitches and RNA thermometers are regulatory sequences that are part of transcripts and influence the translation based on the concentration of selected small molecules and temperature change, respectively. For the prediction of these riboswitches and RNA thermometers, ANNOgesic searches [48] the sequences which are between TSSs (or starting point of a transcript if no TSS was detected) and downstream CDSs, as well as associated with ribosome binding site in the Rfam database using Infernal [49].

### 3.9 Comparison between ANNOgesic predictions and published databases for *Escherichia coli* K12

In order to assess the performance of ANNOgesic, we compared its predictions based on a dRNA-Seq data set and conventional RNA-Seq of *Escherichia coli* K12 MG1655 by Thomason *et al*. [65] and McClure *et al*. [42] with the entries in several databases [20,22,24,31,41,51]. Most of the benchmarking features can be precisely detected (80% or more) (Table S2). Moreover, the predicted features not found in published databases have high possibility to be novel features which are strongly supported by RNA-Seq data (Figure S7B and S7D). TSSs represent an exception with lower success rates and we assume this is mostly due to the higher sensitivity of the dRNA-Seq method in comparison to older protocols. To test this assumption and to investigate the quality of the TSS entries in RegulonDB, we compared the three deposited TSS data sets (Salgado *et al*. generated with Illumina RNA-Seq as well as Mendoza-Vargas *et al*. generated with Roche 454 high-throughput pyrosequencing and generated with Roche 5‘RACE [43,53]) to each other and found very small overlap (Figure S15). Additionally, the 50 nucleotides upstream of TSSs were extracted and scanned with MEME [2] for common motifs that are similar to the ones described for promoters. Only for a small number – 0 to 7% – of TSSs such motifs were found (Table S3) while 80% of the TSS predictions from ANNOgesic have such a promoter motif located upstream (Figure S6C). The same analysis could not be performed with EcoCyc [31] which is lacking TSS information and provide only positions but no strand information for promoters. Due to these results we doubt that the data in those data bases represents a solid ground for benchmarking the accuracy of ANNOgesic’s TSS predictions.

## 4 Discussion

While RNA-Seq has become a powerful method to annotate genomes, the integration of the data is usually very laborious and time-consuming. It requires bioinformatic expertise and involves the application of different programs to perform the different required steps. Here we presented ANNOgesic, a modular, user-friendly annotation tool for the analysis of bacterial RNA-Seq data that integrated several tools, optimizes their parameters, and includes novel prediction methods for several genomic features. With the help of this command-line tool, RNA-Seq data can be efficiently used to generate high-resolution annotations of bacterial genomes with very little manual effort. Besides the annotation files in standard formats, it also returns numerous statistics and visualizations that help the user to explore and to evaluate the results. While it ideally integates conventional RNA-Seq (beneficial for detecting 3’-ends of transcripts) as well as dRNA-Seq (required for the efficiently detection of internal TSSs) as input together (see Figure S16 and S17), it can also perform sufficient predictions with only one class of data for the majority of the genomic features (Table S2).

The performance of ANNOgesic has been here demonstrated by applying it on two published data sets and comparing the results to manually-conducted annotations. ANNOgesic could detect 90% and 83% of the manually-annotated sRNAs *H. pylori* 26695 and *C. jejuni* 81116, respectively. The sRNAs missed by ANNOgesic can be explained by low coverage, not being associated with TSSs, or lack of expression in the assayed conditions (see Figure S18 and S19).

Besides the analyses presented as examples in this study (*H. pylori* 26695 and *C. jejuni* 81116), ANNOgesic was meanwhile successfully applied for detecting transcripts, sRNAs, and TSSs in additional annotation projects (e.g. *Pseudomonas aeruginosa* [14] and *Rhodobacter sphaeroides* [52]. Despite the fact that the program was developed mainly with a focus on bacterial genomes, it has also been used to annotate archaeal genomes (namely *Methanosarcina mazei* (Lutz *et al*., unpublished)) and eukaryotic genomes which have no introns (*Trypanosoma brucei* (Müller *et al*., unpublished)).

ANNOgesic is freely available under an OSI compliant open source license (ISCL) and an extensive documentation has been generated in order to guide the novice and advanced users.

## 5 Conclusion

ANNOgesic is a powful tool for annotating numerous available genome annotations based on RNA-Seq data from multiple protocols. ANNOgesic not only integrated the available tools, but also improve their performance by optimizing parameters, and removing false positives. For the genomic features which can not be detected by currently available tools several novel methods were developed and implemented as part of ANNOgesic. A comprehensive documentation and useful statistics as well as visualization are also provided by ANNOgesic.

## 6 Availability of supporting source code and requirements

- Project name: ANNNOgesic
- Project home page: GitHub - https://github.com/Sung-Huan/ANNOgesic. PyPI - https://pypi.org/project/ANNOgesic/. DockerHub - https://hub.docker.com/r/silasysh/annogesic/.
- Operating system(s): Linux, Mac OS
- Programming language: Python
- Other requirements: Please check the documentation (http://annogesic.readthedocs.io/en/latest/required.html).
- License: ISC (Internet Systems Consortium license simplified BSD license).

## 7 Availability of supporting data

Shell scripts that implement the data analysis workflows as well as the results of the analyses were deposited in Zenodo (https://doi.org/10.5281/zenodo.1161115).

## 8 Additional files

Figure S1: Workflow charts of ANNogesic modules.

Figure S2: Distribution of TSS classes.

Figure S3: Concept and examples for detecting coverage decrease of terminators.

Figure S4: Terminator prediction approach based on convergent genes.

Figure S5: Length distribution of UTRs.

Figure S6: The promoter motifs detected in *Helicobacter pylori* 26695, *Campylobacter jejuni* 81116, and *Escherichia coli* K12 MG1655.

Figure S7: Examples of known and novel intergenic sRNAs that ANNOgesic can detect.

Figure S8: Examples of detected antisense and UTR derived sRNAs.

Figure S9: The coverage plots of the benchmarking sRNA HPnc4620.

Figure S10: Histograms of ranking number of the sRNA benchmarking set.

Figure S11: The distributions of GO term.

Figure S12: The distributions of subcellular localizations.

Figure S13: Visualization of protein-protein interactions.

Figure S14: The example of CRISPR in *Campylobacter jejuni* 81116.

Figure S15: The overlap of three previously published TSS datasets in RegulonDB.

Figure S16: The predicted sRNA which can be detected only in data RNA-Seq after transcript fragmentation.

Figure S17: The comparison between dRNA-Seq and RNA-Seq after transcript fragmentation for detecting transcript

Figure S18: The lowly expressed sRNA - HPnc4610.

Figure S19: An example of known sRNA – CJnc230 – which is not associated with a TSS.

Equation S1: The ranking system of sRNA prediction.

Table S1: The novelties and improvements of genomic feature detection in ANNOgesic.

Table S2: The comparison between ANNOgesic predictions and several databases.

Table S3: The number of TSSs and their associated promoter motifs in RegulonDB.

Table S4: The manual-curated TSS set of *Helicobacter pylori* 26695 (1-400bp).

Table S5: The manual-curated TSS set of *Campylobacter jejuni* 81116 (1-400bp).

Table S6: The manual-curated PS set of *Helicobacter pylori* 26695 (1-400bp).

Table S7: The manual-curated PS set of *Campylobacter jejuni* 81116 (1-400bp).

## 9 Abbreviations

- TSS: transcriptional start site
- PS: processing site
- sRNA: small RNA
- sORF: small open reading frame
- dRNA-Seq: differential RNA Sequencing
- kb: kilo base
- CDS: coding sequence
- CRISPR: clustered regularly interspaced short palindromic repeat
- bp: base pair
- nt: nucleotide
- GO: Gene Ontology

## 10 Competing interests

The authors declare that they have no competing interests.

## 11 Funding

The project was funded by the German Research Foundation (DFG) as part of the Transregio 34 (CRC-TRR34).

## 12 Acknowledgements

We would like to thank several colleagues for fruitful discussion, especially Sarah Svensson, Lars Barquist and Thorsten Bischler for comments regarding the manuscript, Diarmaid Tobin and Till Sauerwein for giving feedback regarding code and documentation.

